# **riboviz**: analysis and visualization of ribosome profiling datasets

**DOI:** 10.1101/100032

**Authors:** Oana Carja, Tongji Xing, Joshua B. Plotkin, Premal Shah

## Abstract

Using high-throughput sequencing to monitor translation in vivo, ribosome profiling can provide critical insights into the dynamics and regulation of protein synthesis in a cell. Since its introduction in 2009, this technique has played a key role in driving biological discovery, and yet it requires a rigorous computational toolkit for widespread adoption. We developed a processing pipeline and browser-based visualization, **riboviz**, that allows convenient exploration and analysis of riboseq datasets. In implementation, **riboviz** consists of a comprehensive and flexible backend analysis pipeline that allows the user to analyze their private unpublished dataset, along with a web application for comparison with previously published public datasets.

**Availability and implementation:** JavaScript and R source code and extra documentation are freely available from https://github.com/shahpr/RiboViz, while the web-application is live at www.riboviz.org.

## Introduction

Analyses of mRNA abundances have been used to gain insight into almost every area of modern biology (1). But ultimately, it is protein synthesis that is the central purpose of mRNAs in a cell. Although mRNA abundance has been used as a proxy for protein production, the correlation between mRNA and protein levels is weak, likely due to post-transcriptional regulation (2; 3; 4). Ribosome profiling (**riboseq**) now provides a direct method to quantify translation, the next obvious step following quantification of transcript abundances (5; 6). This technique rests on the fact that a ribosome translating a fragment of mRNA protects around 30 nucleotides of the mRNA from nuclease activity. High-throughput sequencing of these ribosome protected fragments (called ribosome footprints) offers a precise record of the number and location of the ribosomes at the time at which translation is stopped. Mapping the position of the ribosome-protected fragments indicates the translated regions within the transcriptome. Ribosomes spend different periods of time at different positions, leading to variation in the footprint density along mRNA transcripts. These data provide an estimate of how much protein is being produced from each mRNA (5; 6). Importantly, ribosome profiling is as precise and detailed as RNA sequencing. Even in a short time since its introduction in 2009, ribosome profiling has played a key role in driving biological discovery (13; 14; 15; 16; 17; 18; 19; 20; 21; 12; 22).

However, ribosome profiling is not without its limitations. In mammalian cells, there can be over 10 million unique footprints. The quantification and processing of these footprints remains a challenge and requires computational and domain-specific knowledge. Despite the similarity of ribosome profiling to RNA-seq, for which there is a well established set of bioinformatic tools, ribosome profiling data differs in how it is distributed across the genome. The map and density of ribosomal footprints across the genome contain much more additional information that RNA-seq alone, and traditional bioinformatics pipelines are not designed to handle such data.

We developed a bioinformatic toolkit, **riboviz**, for analyzing and visualizing ribosome profiling data, an important step for this technology to reach a broad audience of practicing biologists. In implementation, **riboviz** consists of a comprehensive and flexible backend analysis pipeline along with a web application for visualization.

## Methods

### Mapping and parsing riboseq datasets

A major challenge in analyses of ribosome-profiling datasets is mapping of individual footprints to ribosomal A, P and E site codons. While several ad hoc rules have been developed to assign reads to particular codons based on the read lengths, these rules are not implemented consistently across studies and as a result, comparing foot-printing reads on a gene across datasets remains a challenge. Using a combination of existing tools used for trimming and mapping reads such as *cutadapt* and *bowtie*, and in-house perl scripts, we have developed a simple set of instructions on mapping reads. We have used this pipeline to remap both RNA-seq and foot-printing datasets from published yeast studies to allow comparison of reads mapped to individual genes across different conditions and labs. In addition, researchers can download individual datasets in a flexible hierarchical data format (HDF5) and gene-specific estimates in flat *.tsv* files. The code and documentation for this pipeline are hosted on Github, with a public bug tracker and community contribution (https://github.com/shahpr/RiboViz).

**Figure 1:**
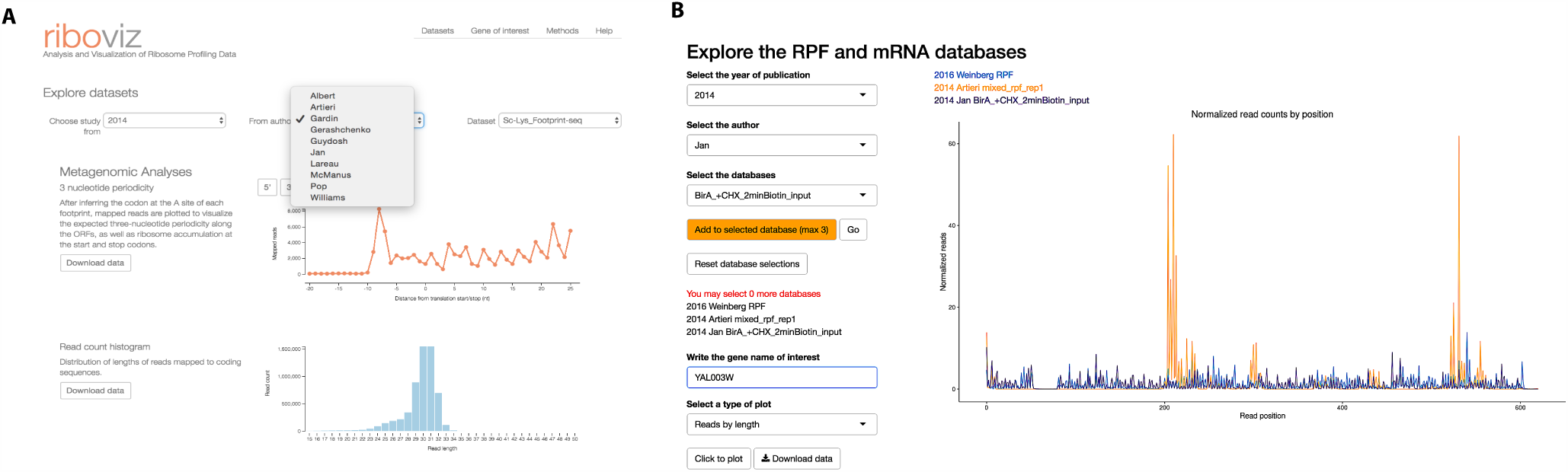
**A.** The **riboviz** website with the user interface allowing dataset selection. **B.** Distribution of reads mapped to YAL003W in three **riboseq** datasets using a Shiny web server.

### Web application and visualization toolkit

The web application is available at www.riboviz.org. Through this web framework, the user can interactively explore publicly available ribosome profiling datasets using JavaScript/D3 (31), JQuery (http://jquery.com) and Boostrap (http://getbootstrap.com) for metagenomic analyses and R/Shiny for gene-specific analyses. The visualization framework of **riboviz** allows the user to select from available riboseq datasets and visualize different aspects of the data. Researchers can also download a local version of the Shiny application to analyze their private unpublished dataset alongside other published datasets available through the **riboviz** website.

**riboviz** allows visualization of metagenomic analyses of (i) the expected three-nucleotide periodicity in footprinting data (but not RNA-seq data) along the ORFs as well as ribosome accumulation of ribosomal footprints at the start and stop codons, (ii) the distribution of mapped read lengths to identify changes in frequencies of ribosomal conformations with treatments, (iii) position-specific distribution of mapped reads along the ORF lengths, and (iv) the position-specific nucleotide frequencies of mapped reads to identify potential biases during library preparation and sequencing. **riboviz** also shows the correlation between normalized reads mapped to genes (in reads per kilobase per million RPKM) and their sequence-based features such as their ORF lengths, mRNA folding energies, number of upstream ATG codons, lengths of 5′ UTRs, GC content of UTRs and lengths of poly-A tails. Researchers can explore the data interactively and download both the whole-genome and summary datasets used to generate each figure.

In addition to the metagenomic analyses, the R/Shiny integration allows researchers to analyze both foot-printing and RNA-seq reads mapped to specific genes of interest, across different datasets and conditions. The Shiny application allows researchers to visualize reads mapped to a given gene across three datasets to compare (i) the distribution of reads of specific lengths along the ORF, (ii) the distribution of lengths of reads mapped to that gene as well as (iii) the overall abundance of that gene relative to its abundance in a curated set of wild-type datasets.

## Summary

Ribosome profiling has been used in viruses, bacteria, yeast, mice, plants, and human cells. This new technique requires a rigorous computational toolkit to reach full promise. **riboviz** will increase the accessibility of this new technology, increase research reproducibility, and offer an broadly useful toolkit for both the community of systems biologists who study genome-wide ribosome profiling data and also to research groups focused on individual genes of interest. By developing and distributing this sets of tools we hope to remove the need for custom script generation by independent researchers, and to increase the pace of genomics research.

## Funding

This work has been supported by start-up funds from Human Genetics Institute of New Jersey and Rutgers University awarded to PS and a Penn Institute for Biomedical Informatics grant to OC. JBP acknowledges funding from the David & Lucille Packard foundation, Army Research Office, and DARPA.

